# Processing UMI Datasets at High Accuracy and Efficiency with the Sentieon ctDNA Analysis Pipeline

**DOI:** 10.1101/2022.06.03.494742

**Authors:** Jinnan Hu, Cai Jiang, Yu S. Huang, Haodong Chen, Hanying Feng, Donald Freed, Yan Qu, Rui Fan, Zhencheng Su, Weizhi Chen

## Abstract

Liquid biopsy enables identification of low allele frequency (AF) tumor variants and novel clinical applications such as minimum residual disease (MRD) monitoring. However, challenges remain, primarily due to limited sample volume and low read count of low-AF variants. Because of the low AFs, some clinically significant variants are difficult to distinguish from errors introduced by PCR amplification and sequencing. Unique Molecular Identifiers (UMIs) have been developed to further reduce base error rates and improve the variant calling accuracy, which enables better discrimination between background errors and real somatic variants. While multiple UMI-aware ctDNA analysis pipelines have been published and adopted, their accuracy and runtime efficiency could be improved.

In this study, we present the Sentieon ctDNA pipeline, a fast and accurate solution for small somatic variant calling from ctDNA sequencing data. The pipeline consists of four core modules: alignment, consensus generation, variant calling, and variant filtering. We benchmarked the ctDNA pipeline using both simulated and real datasets, and found that the Sentieon ctDNA pipeline is more accurate than alternatives.

## Background

It is well established that tumor cells release fragments of DNA, so-called circulating tumor DNA (ctDNA), into the circulatory system during apoptosis or necrosis^1^. ctDNA fragments retain somatic variants from the original tumor tissue. Its existence and fragmentation pattern in blood can be used to infer the presence of a tumor and its stage^2,3^. Accordingly, ctDNA detection and monitoring can serve as a biomarker target for a number of clinical oncology assays, including treatment selection, minimum residual disease (MRD) monitoring, and early cancer detection^4,5^. Evaluations of somatic variants from ctDNA and other liquid biopsies provide multiple advantages over direct biopsy of tumor tissue, as sample collection is significantly faster and less invasive^6^, and the identified variants are likely to have a better representation across heterogeneous tumors^7^. In addition, for MRD monitoring and early cancer detection, no tumor tissue is available and liquid biopsy may be the only choice.

The rapidly decreasing price and increasing throughput of next-generation sequencing technologies has enabled the development of assays to directly sequence ctDNA. In particular, the high-throughput short-read sequencing is ideal for high-depth multi-gene panels that aim to detect variants of low allele frequencies (AFs). As a result, the adoption of NGS ctDNA assays have soared in precision oncology applications^8,9^. However, challenges remain due to limited sample volume and the number of DNA material required for reliable detection of ultra-low AF variants. For example, a single tube of blood typically yields less than 4mL of plasma, from which less than 30ng of DNA or about 9000 haplotype genome equivalents (hGEs) can be extracted. Due to material loss in the ensuing steps of target capture and library construction, number of hGEs available for sequencing is further down to less than 6000, which is considered as the maximum unique coverage, even at saturation sequencing depths.

While overall base accuracy is high, the most widely used short-read sequencing platform has an error rate of around 1 in 1000 bases, approximating the AFs lower end of clinically significant somatic variants. As a solution, Unique Molecular Identifiers (UMIs) have been integrated into ctDNA assays to further reduce base error rate to more accurately discriminate between background errors and real somatic variants^10^. A UMI tag is usually a short sequence of nucleotide bases, added to one or both ends of a template DNA fragment. The uniquely-tagged DNA fragments are then PCR-amplified and sequenced multiple times. Due to the large number of nucleotide combinations, UMIs attached to DNA fragments on a local genomic region are very likely different from each other. Post-sequencing, a consensus sequence is derived by intelligently collapsing reads with the same UMI, thereby mitigating the errors resulting from PCR amplification and sequencing.

A ctDNA NGS analysis pipeline consists of three key steps: alignment, consensus generation, and variant calling. While BWA-mem is adopted as mainstream aligner for short reads, agreement for best practice has not yet been reached for the other two steps. Several pipelines integrate consensus generation and variant calling within one module, such as “DeepSNVMiner”^11^, “MAGERI”^12^, “smCounter”^13^, and “smCounter2”^14^. Other pipelines adopt separate consensus generation and variant calling tools which may allow for more flexibility and faster adaptation to new data types. Popular consensus generation tools include “Fgbio”^15^ and “Gencore”^16^. Gencore is described as a consensus-based dedup tool that works with or without UMIs, while providing higher accuracy compared to alternatives. Popular somatic variant callers include “Mutect2”^17^, “VarDict”^18^, “Strelka2”^19^, and “VarScan2”^20^. Although none of these callers were originally designed for ctDNA assays, they have all demonstrated reasonable accuracy in recent benchmark studies^14,21^.

The NIST-lead Genome in a Bottle (GIAB) consortium has made great advances in providing reliable reference materials and benchmark datasets for germline variant calling^22^, with the first of these datasets available in 2014. However, somatic variant benchmarking has lagged due to the lack of publicly available reference materials and datasets. This lack of somatic reference materials was recently addressed by the Sequencing Quality Control2 (SEQC2) project, which published in 2021 the first multi-center comprehensive project that provided reference datasets and truth sets for both tissue^23,24^ and ctDNA samples^21^. In the SEQC2 ctDNA benchmark study, 360 datasets from two reference samples were generated, providing valuable resources for the whole community.

In this study, we present the Sentieon ctDNA pipeline, a fast and accurate solution for small somatic variant calling from ctDNA sequencing data. The pipeline consists of four core modules: 1) Sentieon BWA, an accelerated version of BWA-mem for alignment^25^; 2) Sentieon Consensus, a consensus generation tool for grouping and generating consensus reads with and without UMI tags; 3) TNscope, a haplotype-based somatic variant caller with high sensitivity^26^; 4) TNscope-filter, a customizable filtering tool for removing false positive variants. We benchmarked the ctDNA pipeline using both simulated and real datasets, and found that the Sentieon ctDNA pipeline is more accurate and faster than alternatives.

## Results

We used four datasets to benchmark accuracy of the pipeline, including a simulated dataset, two in-vitro mixtures with known ground truth, and a real-world dataset of clinical MRD samples.

### Simulated Dataset

UMI-aware consensus generation is considered as a key analysis step for high-depth liquid biopsy sequencing, as downstream variant calling relies on the consensus reads to accurately model base quality in the presence of base errors. In the first benchmark, we generated reads with simulated errors to test the consensus module, comparing the results produced by the Sentieon Consensus tool to Fgbio’s CallMolecularConsensusReads.

To fully simulate the library construction and sequencing process, we initially started with 5 million virtual DNA fragments randomly generated from the human reference genome, with lengths ranging from 100 to 500bp. We then conducted simulated UMI-tagging, PCR amplification and sequencing of these fragments. In total, 8 PCR cycles were simulated and SNP/Indel errors were introduced during the PCR process at assigned error rate, creating a total of 256 virtual sequences for each fragment. Three of the 256 sequences were randomly chosen for use in the simulated sequencing process. During the simulated sequencing, sequencing errors were injected by ART tool using HiSeq2500 error model. In total, 15 million reads with length from 30 to 250bp were generated.

These reads were processed by both the Sentieon Consensus tool and Fgbio to generate consensus reads, which were compared with the ground truth sequence (the initial sequence prior to the introduction of library prep and sequencing error) to evaluate the consensus calling accuracy. The entire simulation, consensus generation, and evaluation process is illustrated in Fig 2.

**Figure 1.**
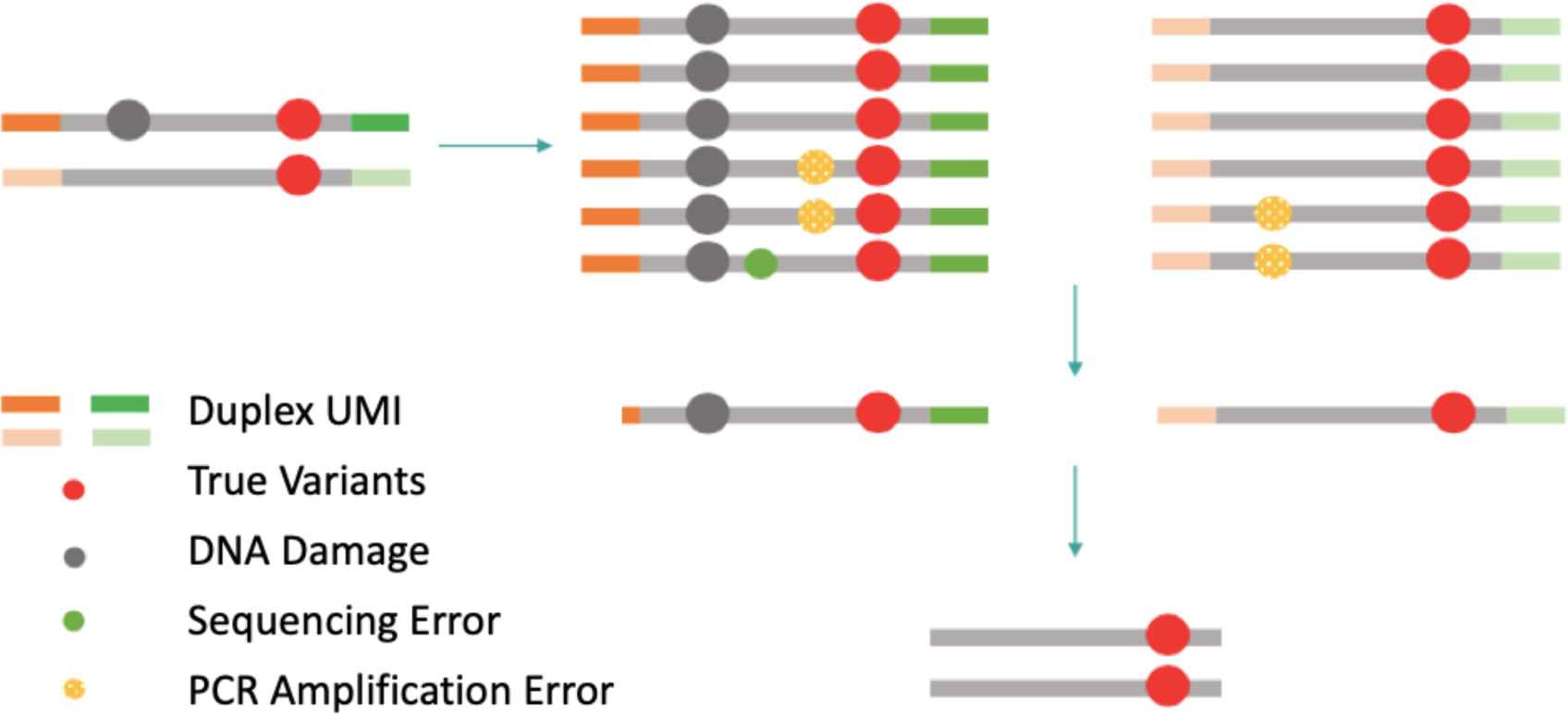
Schematic diagram of duplex UMI error correction. Reads with the same start and end positions, and the same UMI tag are grouped and collapsed into a single consensus read. Relative to standard UMI library preparations, duplex UMI allows for discrimination of different strand of the input molecule, enabling detection and correction of strand-specific errors in the template molecule.

**Figure 2.**
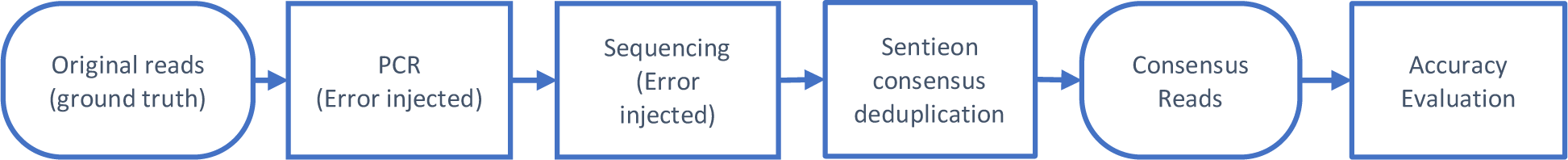
Simulation and benchmarking pipeline.

Three reads in each UMI group were provided to a consensus generation tool, so every consensus base was generated from 3 simulated bases. To better understand the effects of the input bases on consensus accuracy, we assigned each position into one of the categories depending on the input read sequence: three bases are the same (all same, e.g., AAA), two bases are the same (partially same, e.g., AAC), and three bases are all different (all different, e.g., ACG). Over 800 million bases are in the “all same” category, approximately 100 million bases are in the “partially same” category, and very few reads belong to “all different” category. In all three categories, error rate of Sentieon is at least 2 orders of magnitudes lower than Fgbio’s result (Fig 3A).

**Figure 3.**
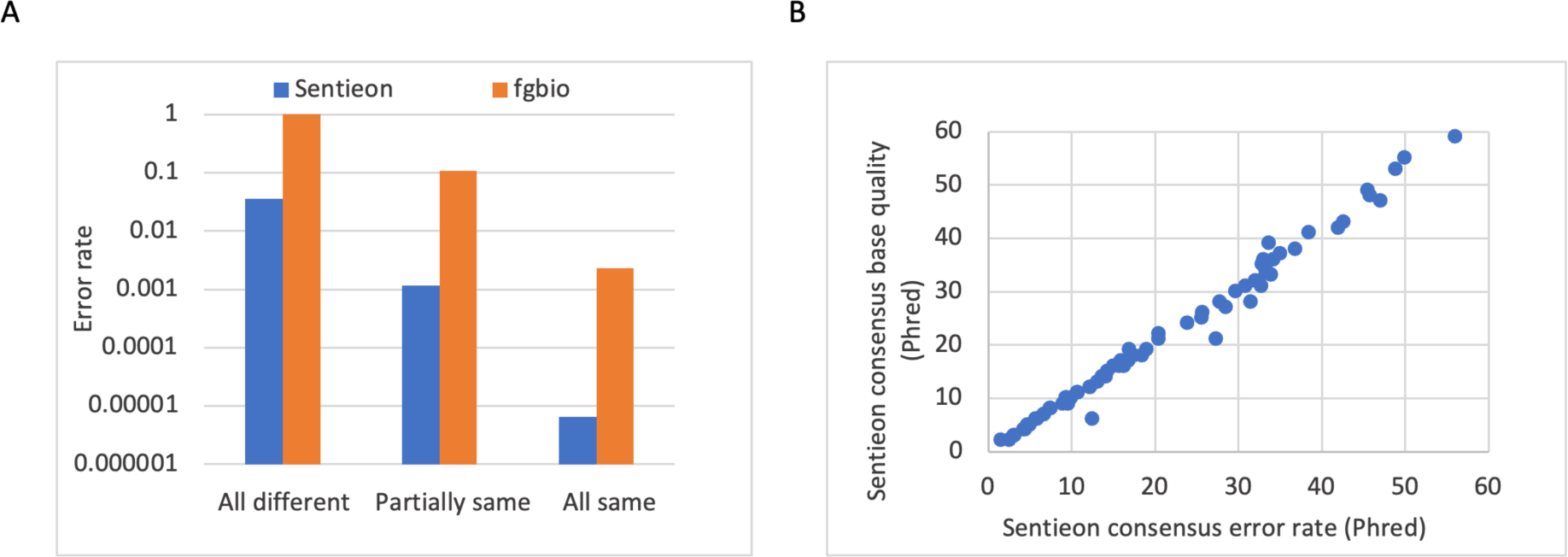
(A) Error rates of Sentieon UMI Consensus tool and the Fgbio. For completely discordant input reads, Fgbio fails to call a consensus base (instead producing an ‘N’ base in the output). (B) The consensus base quality reported by the Sentieon Consensus tool is strongly correlated with the consensus base error rate relative to the ground truth.

The high error rate of Fgbio is likely due to its inaccurate estimation of the PCR amplification error rate, and its treatment of low-quality bases by simply marking them as “N”. This can be easily observed in the “All different” category group where Fgbio fails to return a single correct result, while the Sentieon Consensus tool processes the same input reads with an error rate of under 5%. It should also be noted that in cases where all input bases are the same (the “All same” category), Sentieon and Fgbio still rarely call an incorrect base, as measured comparing the consensus base to the ground truth. In this case, it is likely that all three input reads share an incorrect base, possibly due to an early PCR cycle error.

In addition to correctly generating the consensus sequence, correctly quantifying the confidence in the base as the consensus base quality is also important for downstream variant calling. In theory, the base quality assigned to a consensus base should accurately reflect the true probability of a base being called wrongly. Using the base qualities in the earlier simulation study, we confirm that the Sentieon Consensus tool accurately models the consensus base error rate (Fig 3B).

### In-Vitro Mixture of DNA from Healthy Individuals

Next, we utilized in-vitro mixtures to benchmark the entire ctDNA pipeline, coupling the Sentieon Consensus module with TNscope. Sentieon TNscope is a haplotype-based somatic variant caller that follows the general mathematical models in the GATK Mutect2, with a variety of improvements. To evaluate the accuracy of the Sentieon ctDNA pipeline, we utilized two in-vitro mixture datasets with known variant calls for benchmarking, and compared the accuracy with an alternative consensus generation and variant calling pipeline.

The first dataset was generated using a mixture of extracted DNAs from two healthy individuals (Fig 4). Truthset variants for each individual were determined by the GATK germline variant calling pipeline. One individual’s DNA (spike-in) was then mixed with another’s (background) at 0.2% and 0.3% titration rates. In total, three libraries were constructed from mixed DNA samples. A customized panel probe hybridization was conducted on all three libraries, covering 57 homozygous and 70 heterozygous truth SNPs in the panel region. The libraries were then sequenced using Illumina or MGI platforms and the datasets were subsampled to 30,000x before analysis. Detailed info for each dataset is listed in Table 1, and details of the sample collection and processing procedures are described in the method section.

**Figure 4.**
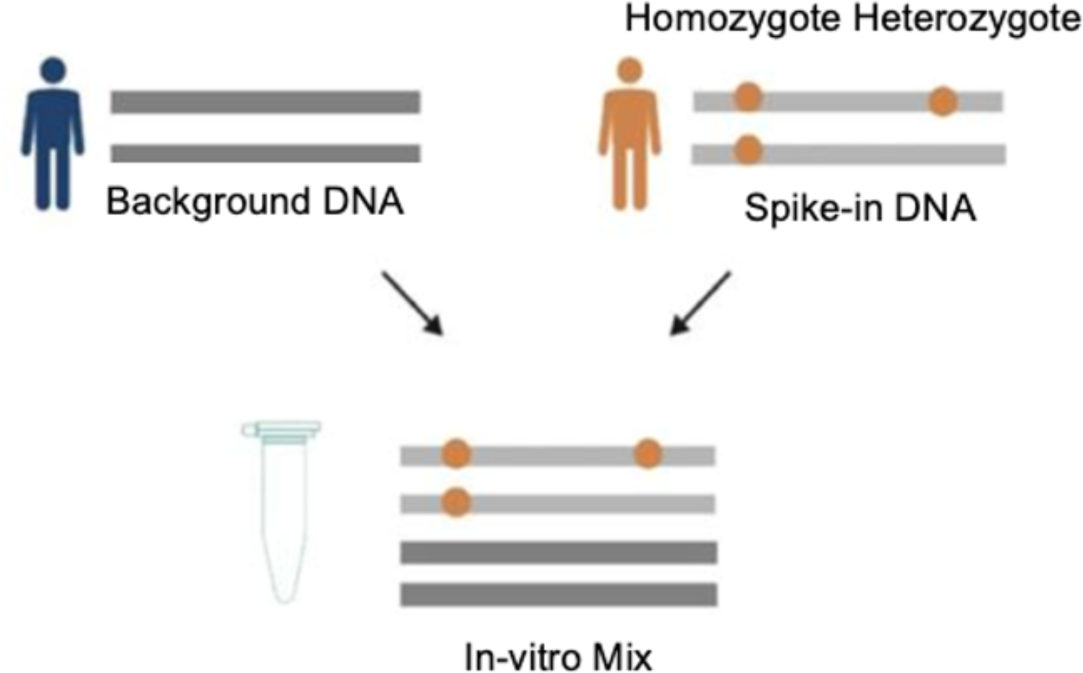
In-Vitro mixture samples prepared by titration of DNAs from two healthy individuals.

**Table 1.**
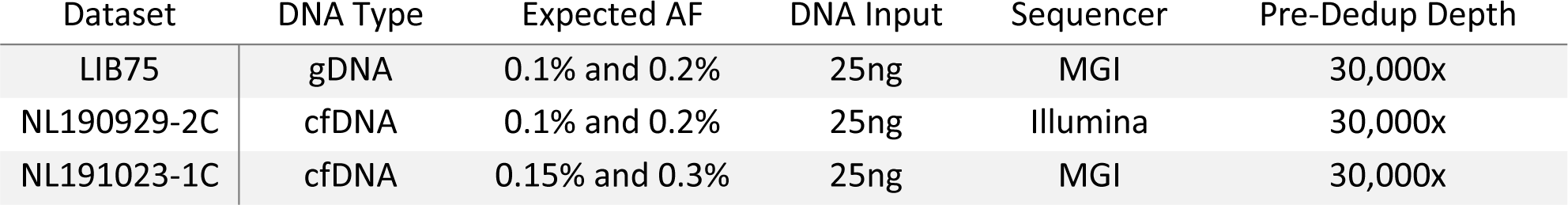
Wet lab settings of the MRD datasets.

The three datasets are analogous to MRD detection, with an expected mutation frequency down to 0.1%. DNA type and sequencing platforms are varied, to test analysis pipeline’s performance on different wet lab settings.

The three datasets were processed by the Sentieon ctDNA pipeline, and an “Fgbio + Vardict” pipeline as the alternative. The resulting variant calls were compared against the truth set for accuracy calculation.

Accuracy metrics for the two pipelines are reported in Fig 5. The Sentieon pipeline returned slightly superior overall F-Score and higher recalls.

**Figure 5.**
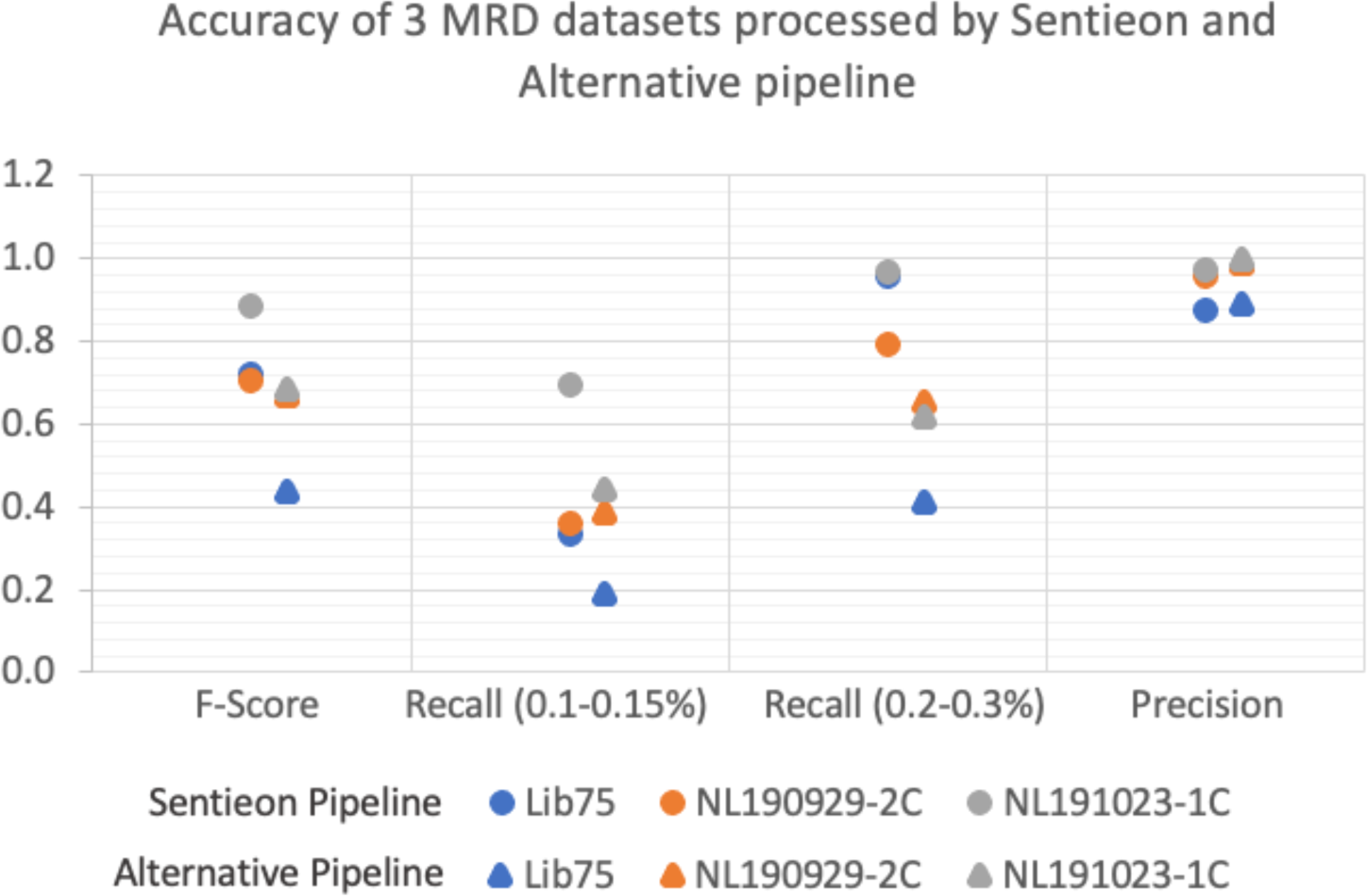
Accuracy of the Sentieon ctDNA pipeline and alternative pipeline, on MRD datasets.

**Figure 6.**
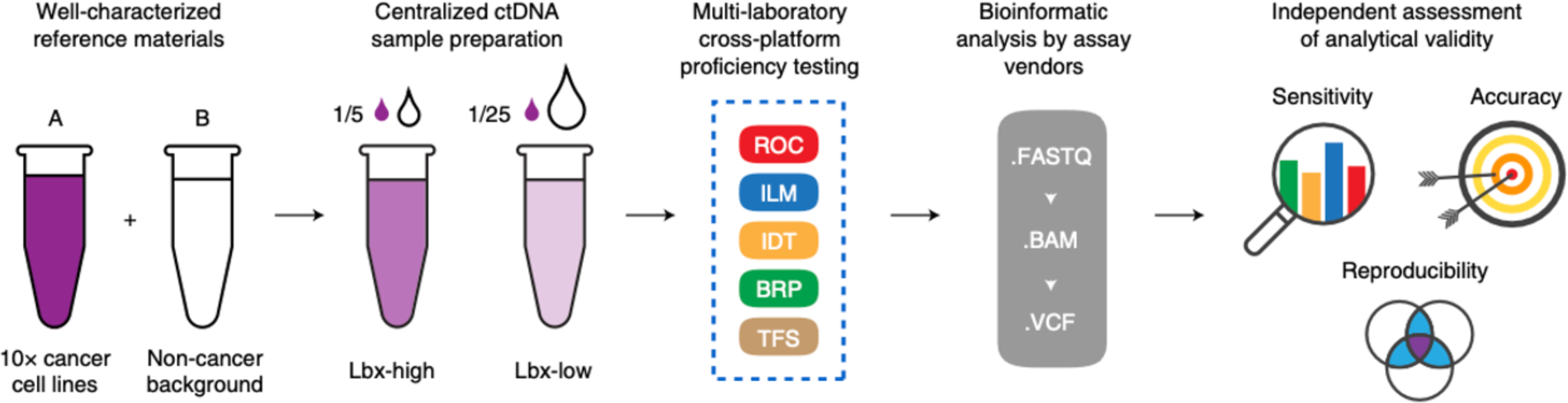
Schematic diagram of reference sample generation, assay process, variant calling, and accuracy benchmark^21^.

### SEQC2 ctDNA Dataset

In addition to the in-vitro mix samples described above, we also utilized the recent published SEQC2 dataset^21^ to benchmark the performance of the Sentieon ctDNA pipeline. The SEQC2 study is a large multi-center project that aims to generate ctDNA reference samples, and to benchmark current available ctDNA assays. Accordingly, the SEQC2 project generated reference samples through in-vitro mixture of two cell line DNA samples with known somatic variants, at different titration rates. The “Lbx-high” mixture contains variants with median frequency around 1%, and majority above 0.5%; while the “Lbx-low” mixture contains variants with median frequency around 0.2% and majority above 0.1%. Both reference DNAs were then sent to multiple ctDNA assay vendors for sequencing and bioinformatics analysis. BRP (Burning Rock Dx) assay provided the highest accuracy in the project, and we therefore chose the BRP datasets for our benchmark. Details of dataset prep methods can be found in the SEQC2 study paper^21^.

We chose to work with eight Lbx-low datasets, because their lower AFs could better highlight the advantage of UMI tagging and consensus analysis. The fastq files for these samples were processed by the Sentieon ctDNA pipeline, and variant calling accuracy was evaluated using the SEQC2 truth set. Performance was compared against the BRP analysis pipeline, which is based on a BRP-developed UMI consensus tool and VarScan2^21^. The depth of each sample is reported in Table 2. The post-dedup depth reflects hGEs being sequenced, and does not have a strong correlation with pre-dedup depth. Average group size for each UMI consensus read is approximately 7 to 12.

**Table 2.** Pre-dedup and post-dedup depth of the evaluated datasets.

| Datasets | Site | Library | Pre-Dedup Depth | Post-Dedup Depth |
| --- | --- | --- | --- | --- |
| SRR13200999 | 25 | 1 | 41,006x | 5,693x |
| SRR13200998 | 25 | 2 | 71,212x | 5,760x |
| SRR13200997 | 25 | 3 | 48,660x | 5,181x |
| SRR13200996 | 25 | 4 | 67,624x | 5,707x |
| SRR13200991 | 26 | 1 | 37,979x | 5,352x |
| SRR13200990 | 26 | 2 | 37,164x | 5,361x |
| SRR13200988 | 26 | 3 | 48,317x | 5,667x |
| SRR13200987 | 26 | 4 | 40,972x | 5,433x |

**Table 3:**
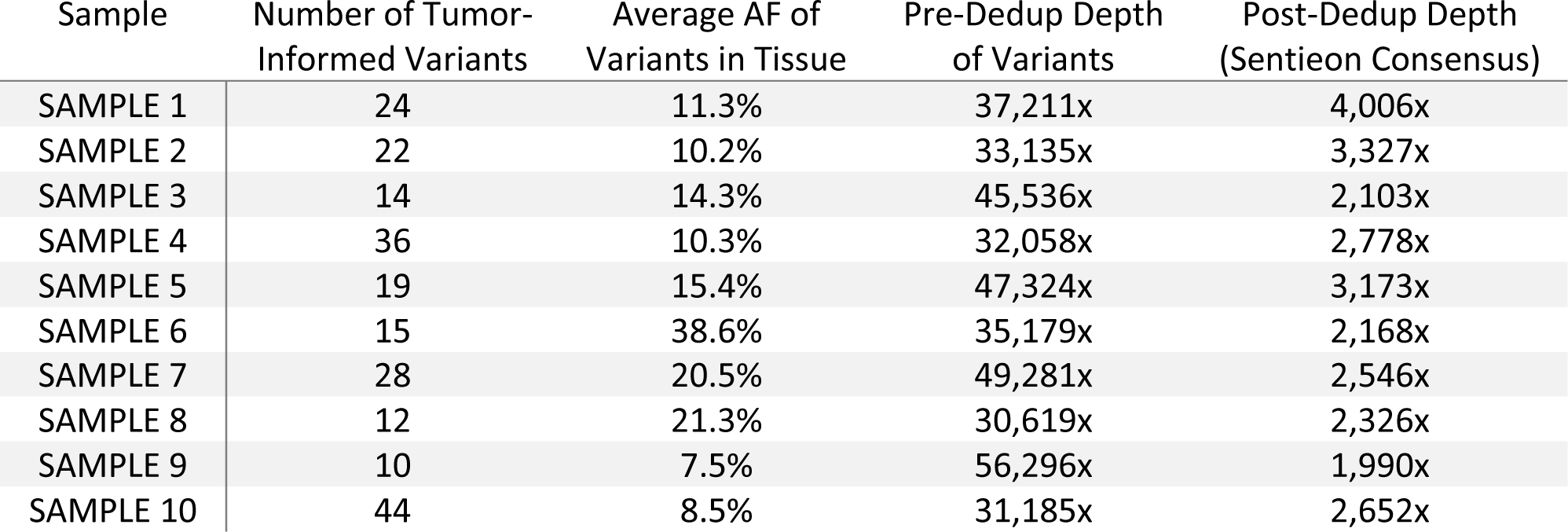
The ten clinical samples contain 224 known somatic mutations in total. Their AFs from original tumor tissues, along with the pre and post consensus depths are shown.

| Sample | Number of Tumor-Informed Variants | Average AF of Variants in Tissue | Pre-Dedup Depth of Variants | Post-Dedup Depth (Sentieon Consensus) |
| --- | --- | --- | --- | --- |
| SAMPLE 1 | 24 | 11.3% | 37,211x | 4,006x |
| SAMPLE 2 | 22 | 10.2% | 33,135x | 3,327x |
| SAMPLE 3 | 14 | 14.3% | 45,536x | 2,103x |
| SAMPLE 4 | 36 | 10.3% | 32,058x | 2,778x |
| SAMPLE 5 | 19 | 15.4% | 47,324x | 3,173x |
| SAMPLE 6 | 15 | 38.6% | 35,179x | 2,168x |
| SAMPLE 7 | 28 | 20.5% | 49,281x | 2,546x |
| SAMPLE 8 | 12 | 21.3% | 30,619x | 2,326x |
| SAMPLE 9 | 10 | 7.5% | 56,296x | 1,990x |
| SAMPLE 10 | 44 | 8.5% | 31,185x | 2,652x |

The Precision and Recall for the two pipelines are shown in Fig 7. On the same high-depth UMI dataset, the Sentieon pipeline outperformed the BRP analysis pipeline, in terms of both Recall and F1 Score.

**Figure 7.**
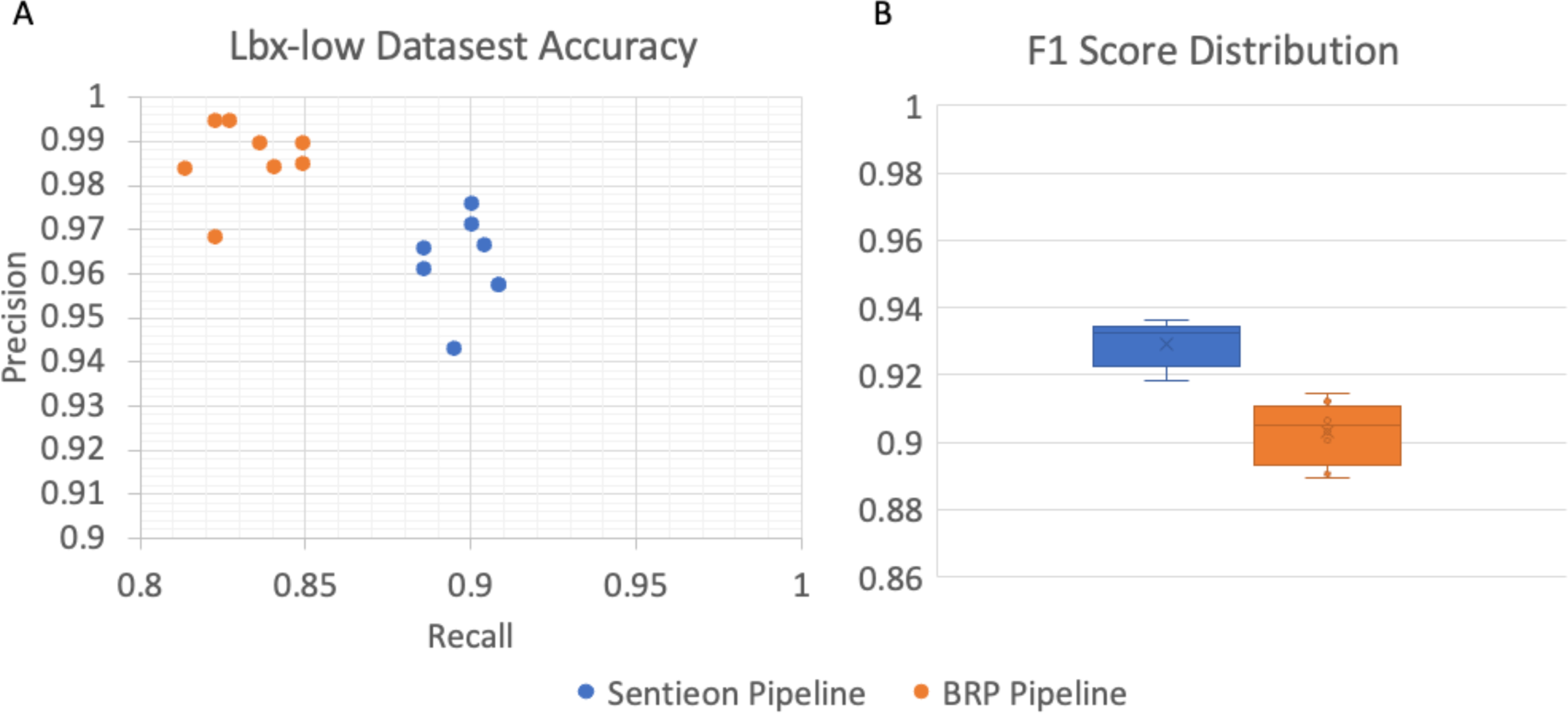
(A) Precision vs. Recall of the 8 Lbx-low datasets; (B) F1 Score Distributions of the two pipeline.

### Clinical Minimum Residual Disease Validation

The prior analyses demonstrate that the Sentieon ctDNA pipeline has good performance on both synthetic datasets as well as in-vitro mixtures. To further assess performance of the pipeline, we tested our methods on real-world clinical samples. The dataset includes 10 clinical samples from a tumor-informed MRD assay. Each sample has a pre-determined set of oncogenic clonal mutations, which were identified through tumor biopsy sequencing for the same patient. The subsequent ctDNA test would only assess the pre-determined mutation set to call if a variant is positive or negative, serving as indicator of patient’s disease status. By tracking only the pre-determined set of clonal mutations, the tumor-informed MRD approach eliminates the potential interference from other non-tumor (i.e. clonal hematopoiesis) somatic variants. Thus, the limit of detection (LOD) of tumor-informed MDR approaches can reach to below 0.1%, which is much lower than tumor-naïve approaches.

First, reads of these 10 samples were aligned using Sentieon BWA to generate pre-dedup BAM files. After alignment, three UMI consensus generation modules including Genecast MinerVa, Sentieon, and Fgbio were respectively applied. Given all reads aligned to a same genomic position, the MinerVa UMI module clusters them into different clusters under the rule that all UMIs within a cluster are within one edit distance of each other, and then derives the consensus read for each cluster using a statistical model that considers the base qualities of each read. MinerVa UMI also has the ability to run in a per-variant mode for higher sensitivity and faster speed.

After consensus read generation, pre-determined mutations were examined in the consensus reads for the number of supporting reads and total consensus read depth, and VAF (Variant Allele Frequency) was calculated at each variant site. The Genecast MinerVa MRD pipeline then produces a p-value for each identified variant based on a proprietary statistical model, which is trained on a population dataset constructed from approximately 1000 healthy individuals. For Sentieon and Fgbio, a variant is called positive when its p-value is less than 0.05; while for Genecast MinerVa, the criteria is p-value less than 0.05, or a duplex count greater than 0.

Based on the results (Fig 8), Genecast MinerVa has the highest positive rate (= the number of detected ctDNA variant / the size of the pre-determined variant set), while Sentieon and Fgbio have slightly lower positive rate. Note that the consensus base quality information generated by Sentieon or Fgbio was not used in the variant calling, which suggests room for improvement.

**Figure 8.**
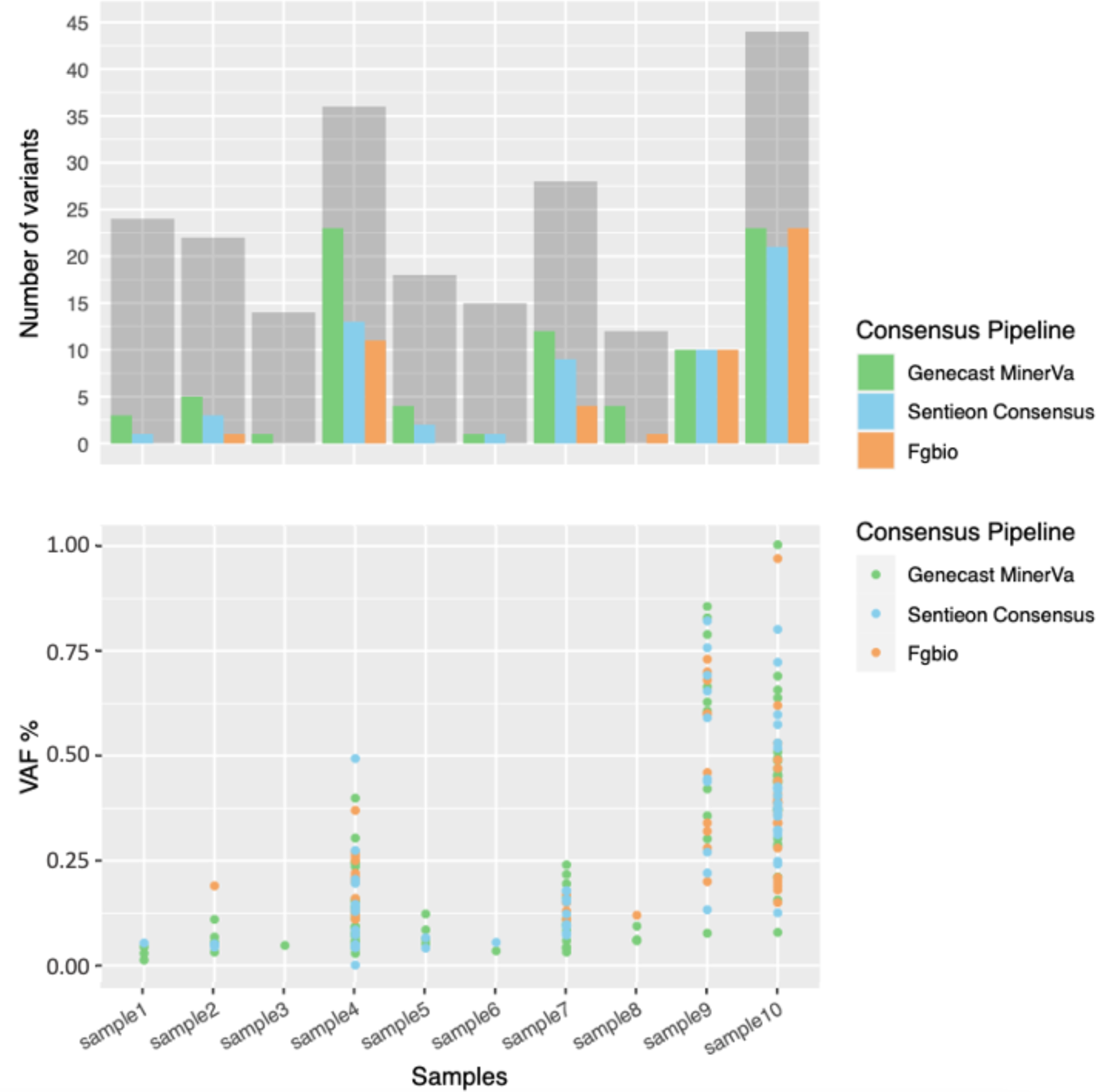
Comparing the ctDNA variant detection from post-dedup BAMs generated by three tools. Upper panel: The height of a gray bar represents the size of the pre-determined variant set from the tumor tissue sequencing. Within each gray bar, a green/blue/orange bar represents the number of ctDNA variants detected in the same sample. Lower panel: The VAFs of all detected ctDNA variants are below 1%, with the majority below 0.25%.

While the true positive rate for clinical samples is hard to know, similar positive rates from different tools corroborate the usefulness of the UMI-consensus generation approach for real-world clinical assays. Furthermore, we assessed the specificity of each tool using a cross-patient scheme. In this scheme, for each patient, non-overlapping somatic mutations from other patients were considered as potential for false positive variant detection, and were combined to form the “negative” pre-determined variant set for tracking. A total of 1998 “negative” variants were assessed in this cross-patient analysis. From these potential false positives, 5 out of 1998 variants were called positive by Genecast MinerVa, 8 by Fgbio, and 10 by Sentieon Consensus. Based on these data, we conclude that the specificity for all three methods is above 99.5%.

### Runtime Comparison

We designed two runtime comparison datasets to benchmark the Sentieon Consensus tool and the whole Sentieon ctDNA pipeline.

The Sentieon Consensus is optimized for efficient processing of UMI-tagged reads while maintaining high accuracy for UMI consensus reads. Improvements in the underlying algorithms and a performance-focused software implementation allow the Sentieon pipeline to process the data ∼20X faster (4,317sec vs 82,679sec) relative to Fgbio (Fig 9A). This benchmark was conducted using a 0.5% AF dataset (N13532 from benchmark study^14^) on a 32-logical-core Intel Xeon platform.

**Figure 9.**
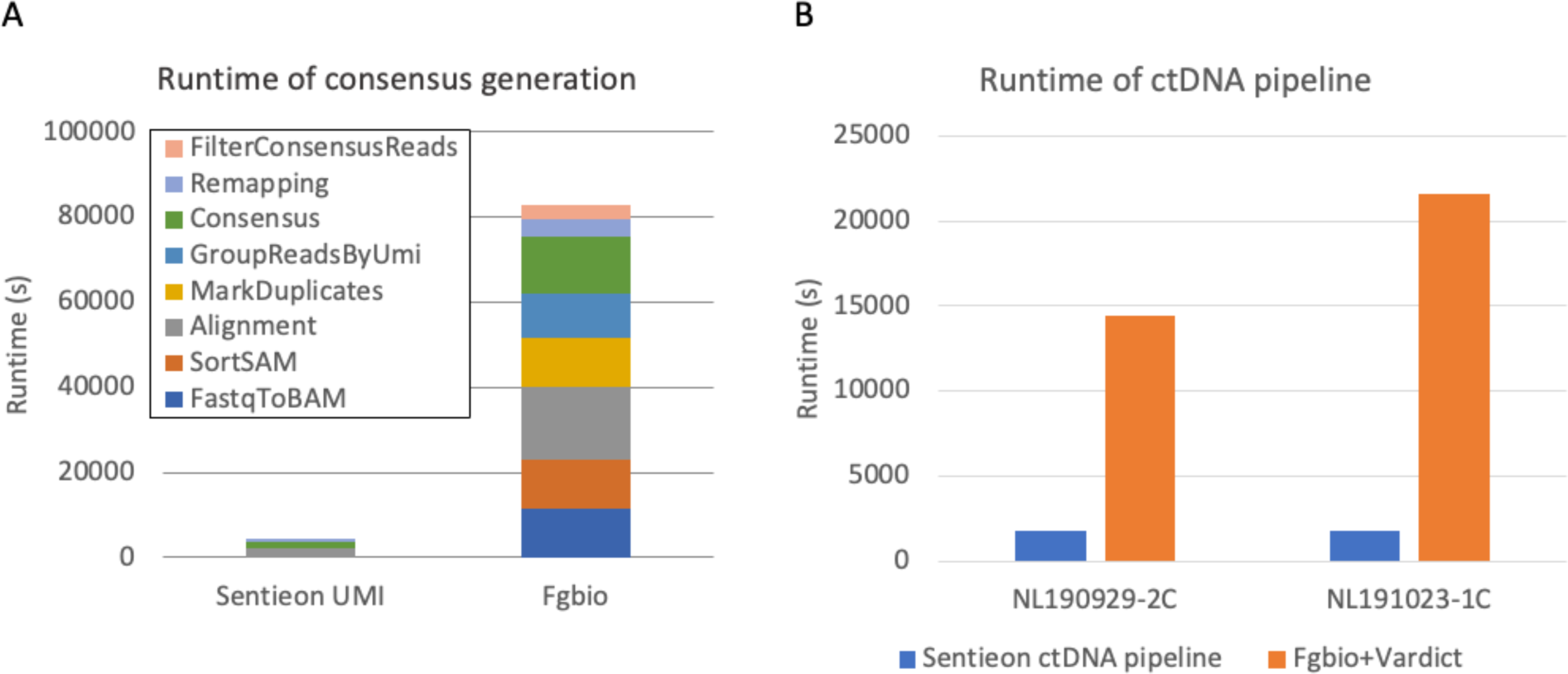
**A**. Runtime comparison of the consensus generation step. The Sentieon UMI Consensus tool showed 20X speed-up over Fgbio. **B**. Runtime comparison of the whole ctDNA processing pipeline including alignment, consensus generation, variant calling and filtering.

The whole pipeline runtime comparison was conducted using two in-vitro mixture datasets at the original sequencing depth. Ten threads were allocated to each pipeline. For the “Fgbio + Vardict” pipeline (alternative pipeline), the dataset was split into 10 chunks and run in parallel to improve its speed. The alternative pipeline is about 10x slower than the Sentieon ctDNA pipeline, even with the additional parallelization (Fig 9B).

## Methods

### Healthy Individuals In-vitro Mix Dataset Generation

Genomic DNAs were extracted from two healthy individuals’ blood samples using the TIANamp Genomic DNA kit (#DP304, TIANGEN BIOTECH). Cell free DNAs were extracted from the same individuals using the Serum/Plasma Circulating DNA Kit (#DP339, TIANGEN BIOTECH).

Genomic DNA or cfDNA from one individual (spike-in) was mixed with another individual’s DNA (background) at 0.2% and 0.3% titration rate, respectively. Library 75 was constructed by NadPrep DNA Library Preparation Kit (for MGI) including Bi-Molecular Identifier (BMI) adapter (#1003821, Nanodigmbio) from 25ng gDNA fragmented by NEBNext dsDNA Fragmentase (#M0348, NEB). Library NL190929-2c was constructed by NadPrep DNA Library Preparation Kit (for Illumina) including UMI adapter (#1003431, Nanodigmbio) from 25ng cfDNA. Library NL190023-1c was constructed by NadPrep DNA Library Preparation Kit (for MGI) including Bi-Molecular Identifier (BMI) adapter (#1003821, Nanodigmbio) from 25ng cfDNA. Following end repair and A-tailing, dilution was performed. 40µL of NadPrep SP Beads were used for library clean up and ligated fragments were amplified with 6-9 cycles using 0.5M index primers mix. Library yields were controlled between 500-1000ng. 60µL of NadPrep SP Beads were used to recycle libraries.

All three libraries were processed by single-tube capture hybridization. Following the Nanodigmbio Hybridization capture protocol (NadPrep Hybrid Capture Reagents, #1005101), each pool of DNA was combined with 5µL of 1mg Cot-1 DNA (Invitrogen) and 2µL 0.2nmol NadPrep NanoBlockers (#1006204 for MGI, #1006101 for Illumina, Nanodigmbio) to prevent cross hybridization and minimize off-target capture. Library and blocker were dried and re-suspended in hybridization buffer and enhancer. Target capture with 38kb “SNP ID” Panel Probes (Nanodigmbio) was performed overnight. Streptavidin M270 (Invitrogen) beads were used to isolate hybridized targets according to Nanodigmbio hybridization capture protocol. Captured DNA fragments were amplified with 13–15 cycles of PCR.

Libraries were then sequenced by either 100bp paired-end runs on MGI-2000 sequencer at Nanodigmbio R&D Center, or 150bp paired-end runs on Illumina XTen sequencer at the Wuxi Genome Center. Germline variants of the two individuals were called by the GATK pipeline, and germline variants unique to a single individual were used as truth set for accuracy calculation.

### Sentieon ctDNA Pipeline - Alignment and Sort

The Sentieon Genomics tool set is a suite of software tools for secondary analysis of next-generation sequence data. The Sentieon pipelines consist of optimized implementations of the mathematical models of the most accurate variant calling pipelines. Improvements in performance are achieved through optimization of the algorithms and improved resource management.

Prior to variant calling, sequence data are aligned to the human reference genome using Sentieon BWA. BWA-MEM is one of the most popular aligners for alignment of next-generation sequencing reads given its accuracy and ability to produce correct alignments at structural variant breakpoints. Sentieon provides an optimized implementation of BWA resulting in a 1.0 to 3.9x speedup^25^, while producing identical alignments. Read coordinate sorting was conducted by Sentieon utility module “sort”.

### Consensus Generation

The Sentieon Consensus module performs sophisticated modeling of the base errors introduced by library construction and sequencing process to improve consensus read accuracy. Reads are grouped based on their positions and UMI tags if available, and then the grouped reads are modeled statistically to account for multiple sources of base error. Potential sources of error include, PCR errors occurring during library construction; single-strand errors introduced prior to UMI tagging; and sequencing errors. Parameters of the statistical model are learned and calibrated directly from the dataset without user input. Consensus reads are then called from the grouped reads using a Bayesian model informed by the learned parameters, and overlapping read pairs are then merged. It should be noted that during the consensus calling no reads are discarded, and all read information is counted in for model calibration. Consensus calling confidence is reflected in the assigned base quality of consensus bases, which helps downstream callers improve variant calling accuracy.

### TNscope and TNscope-filter

TNscope is a haplotype-based variant caller that follows the general principles of the mathematical models first implemented in the GATK HaplotypeCaller and MuTect2. This includes active region detection, assembly of haplotypes from the reference and local read data using a de Bruijn-like graph, pair-HMM for calculation of read-haplotype likelihoods followed by genotype assignment. Like MuTect2, TNscope evaluates the tumor and normal haplotypes jointly if the matching normal sample is available, achieving significantly higher precision for somatic variant detection.

Several improvements have been incorporated into the mathematical model of TNscope to increase its recall and precision. The highly-efficient implementation allows TNscope to choose a lower threshold for triggering active regions, facilitating a more comprehensive evaluation of potential variants.

Furthermore, the detected active regions are typically of higher quality as TNscope uses a statistical model to trigger active regions rather than a fixed cutoff. Local assembly is improved, resulting in more frequent identification of the correct variant haplotype. Genotyping is improved as well due to the adoption of a novel quality score model and various nonparametric statistical tests to eliminate false-positive variants. TNscope also outputs several novel variant annotations that can be used for improved variant filtration. Tumor-only mode is enhanced with a panel of non-matching normal samples. For high-depth targeted sequencing data, downsampling is not required due to TNscope’s computational efficiency, which makes it an ideal haplotype-based variant caller for the detection of rare somatic events in ctDNA samples.

TNscope-filter module is a VCF filtering tool, that could identify false positive variants from raw VCF files based on input parameters. Its functionality is similar to BCFtools^27^ but is more compatible with other Sentieon modules.

## Conclusions

In this study, we present the Sentieon ctDNA analysis pipeline and benchmark its accuracy using both simulated and real datasets. Superior recall and precision compared to alternative pipelines was observed in most testing datasets. The superior performance of the Sentieon pipeline is largely due to the sophisticated statistical model used in the consensus generation tool couple with highly accurate somatic variant calling from Senteion TNscope. Besides being more accurate, the Sentieon ctDNA pipeline is also much faster than alternative pipelines, enabling the timely processing of high-depth large-panel datasets.

## Code Availability

Script of Sentieon UMI pipelines benchmarked in this study can be found on Github page: https://github.com/Sentieon/sentieon-scripts/blob/master/example_pipelines/somatic/TNscope/Somatic_ctDNA_with_UMI.sh; TNscope_filter “-x” preset was set as “ctDNA” or “ctdna_UMI”, “-min_tumor_allele_frac” and “-min_tumor_af” parameters were set based on desired AF cutoff.

